# Bacterial modification of the host glycosaminoglycan heparan sulfate modulates SARS-CoV-2 infectivity

**DOI:** 10.1101/2020.08.17.238444

**Authors:** Cameron Martino, Benjamin P. Kellman, Daniel R. Sandoval, Thomas Mandel Clausen, Clarisse A. Marotz, Se Jin Song, Stephen Wandro, Livia S. Zaramela, Rodolfo Antonio Salido Benítez, Qiyun Zhu, Erick Armingol, Yoshiki Vázquez-Baeza, Daniel McDonald, James T. Sorrentino, Bryn Taylor, Pedro Belda-Ferre, Chenguang Liang, Yujie Zhang, Luca Schifanella, Nichole R. Klatt, Aki S. Havulinna, Pekka Jousilahti, Shi Huang, Niina Haiminen, Laxmi Parida, Ho-Cheol Kim, Austin D. Swafford, Karsten Zengler, Susan Cheng, Michael Inouye, Teemu Niiranen, Mohit Jain, Veikko Salomaa, Jeffrey D. Esko, Nathan E. Lewis, Rob Knight

## Abstract

The human microbiota has a close relationship with human disease and it remodels components of the glycocalyx including heparan sulfate (HS). Studies of the severe acute respiratory syndrome coronavirus (SARS-CoV-2) spike protein receptor binding domain suggest that infection requires binding to HS and angiotensin converting enzyme 2 (ACE2) in a codependent manner. Here, we show that commensal host bacterial communities can modify HS and thereby modulate SARS-CoV-2 spike protein binding and that these communities change with host age and sex. Common human-associated commensal bacteria whose genomes encode HS-modifying enzymes were identified. The prevalence of these bacteria and the expression of key microbial glycosidases in bronchoalveolar lavage fluid (BALF) was lower in adult COVID-19 patients than in healthy controls. The presence of HS-modifying bacteria decreased with age in two large survey datasets, FINRISK 2002 and American Gut, revealing one possible mechanism for the observed increase in COVID-19 susceptibility with age. *In vitro*, bacterial glycosidases from unpurified culture media supernatants fully blocked SARS-CoV-2 spike binding to human H1299 protein lung adenocarcinoma cells. HS-modifying bacteria in human microbial communities may regulate viral adhesion, and loss of these commensals could predispose individuals to infection. Understanding the impact of shifts in microbial community composition and bacterial lyases on SARS-CoV-2 infection may lead to new therapeutics and diagnosis of susceptibility.

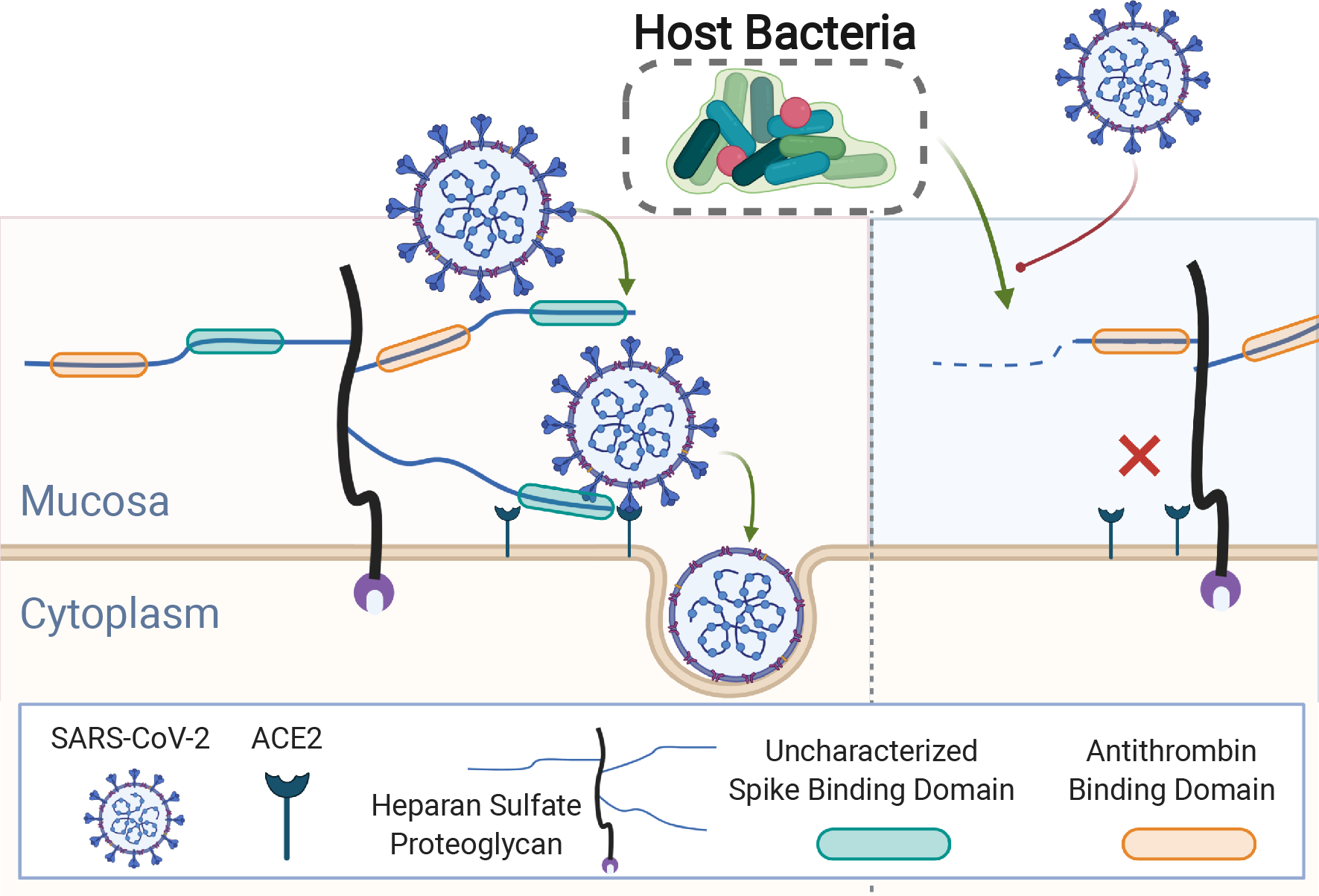
Graphical Abstract. Diagram of hypothesis for bacterial mediation of SARS-CoV-2 infection through heparan sulfate (HS). It is well known that host microbes groom the mucosa where they reside. Recent investigations have shown that HS, a major component of mucosal layers, is necessary for SARS-CoV-2 infection. In this study we examine the impact of microbial modification of HS on viral attachment.

## Introduction

All living cells are coated with a dense forest of glycans and glycoconjugates called the glycocalyx. Many viral pathogens have evolved mechanisms to attach to host glycans in order to bind to tissues and pass through the glycocalyx to engage protein receptors in the plasma membrane required for cell entry (Cagno et al., 2019). Of these, severe acute respiratory syndrome coronavirus (SARS-CoV-2) (Lang et al., 2011), dengue (Chen et al., 1997), human papillomavirus (Giroglou et al., 2001), hepatitis B (Schulze et al., 2007), herpes simplex virus (Spear et al., 1992), human immunodeficiency virus (Connell and Lortat-Jacob, 2013), and others, target heparan sulfate (HS), a highly negatively charged linear polysaccharide present on the surface of all mammalian cells (Esko and Selleck, 2002). Infection by SARS-CoV-2, the causative agent of COVID-19, can be blocked with the HS derivative heparin (Kim et al., 2020; Mycroft-West et al., 2020; Clausen and Sandoval et al., 2020). SARS-CoV-2 attachment and infection requires binding to both HS and angiotensin converting enzyme 2 (ACE2) via distinct regions of the receptor binding domain (RBD) (Clausen and Sandoval et al., 2020). Reducing the interaction between the SARS-CoV-2 spike protein and host-cell HS is therefore an attractive approach to reduce viral docking and subsequent infection.

Several bacterial taxa produce enzymes that modify specific classes of the glycosaminoglycan (GAG) family, including HS, and much of GAG research has been advanced using purified bacterial enzymes (Linhardt et al., 1986). Polysaccharide utilization loci (PUL) are co-localized and co-regulated genes responsible for complex carbohydrate detection and degradation in bacteria (Grondin et al. 2017). PULs have been annotated in *Bacteroides thetaiotaomicron* (human gut isolate; Cartmell et al., 2017; Ndeh et al., 2020; Ulmer et al., 2014) and *Flavobacterium heparinum* (soil isolate; Galliher et al., 1981). These PULs allow the bacteria to degrade and catabolize many complex carbohydrates including HS. The interplay of bacterial and viral utilization of host glycocalyx GAGs provides a mechanism for transkingdom interaction (Pfeiffer and Virgin, 2016) through modulation of pathogen adhesion spanning the eukaryotic host, bacteria in the microbiome, and SARS-CoV-2. Removal of cell-surface HS via heparin lyase (HSase) purified from *F. heparinum* effectively eliminates SARS-CoV-2 virus infection and spike protein binding (Clausen and Sandoval et al., 2020).

Here we show a relationship between SARS-CoV-2 susceptibility and the capacity of the human microbiome to catabolize HS, a critical host factor involved in mediating SARS-CoV-2 infection. A computational model of HS catabolism was developed to assess the capacity of human microbes to catabolize HS. HS-modifying bacteria are reduced in patients with COVID-19. Moreover, the microbial HS-modifying capacity is reduced with age, and enriched in females compared to males, both major COVID-19 susceptibility factors. Finally, evidence shows that common commensal bacteria have the capacity to prevent SARS-CoV-2 spike protein binding by degrading host HS. These results suggest that differences in human microbiome composition may influence SARS-CoV-2 infectivity and help explain the observed sex and age-associated heterogeneity of COVID-19 rates across populations.

## Results

### Common human-associated microbes possess a high capacity for HS modification

Bacteria rely on host glycans for binding, immunological recognition, and as an energy source (e.g. carbon and nitrogen catabolism). For this purpose, bacteria produce a variety of enzymes encoded in PUL capable of degrading the complex glycans making up the host cell glycocalyx. PUL have been annotated in hundreds of bacterial genomes, providing insights into the carbohydrates each species can catabolize (Terrapon et al., 2015, 2018). These activities have also been formulated in a computational modeling platform to simulate PUL activity (Eilam et al., 2014). As a further generalization of PUL estimation beyond comprehensive genome annotation or mechanistic modeling, we used a pathway-based approach by modeling catabolism of each major glycan as a metabolic task in order to predict bacteria involved in catabolism of HS (Richelle et al., 2018, 2019, 2020).

Recent work demonstrated the necessity of HS for SARS-CoV-2 cellular binding and efficient infection (**Fig. 1A**) (Clausen and Sandoval et al., 2020), suggesting that the microbial HS catabolic capacity in the native human gut microbiome might affect infection. We curated a metabolic task (i.e. a group of catabolic reactions necessary to transition between metabolites (Richelle et al., 2020)) that describes HS modification **(Fig. 1B)**. Using this task, we quantified the total counts aligned to each glycosylation-associated gene in each microbe in the FINRISK 2002 dataset, a fecal shotgun metagenomic dataset spanning over 6,000 samples from participants ranging from 25–74 years old (Borodulin et al. 2015; Salosensaari et al., 2020).

**Figure 1.**
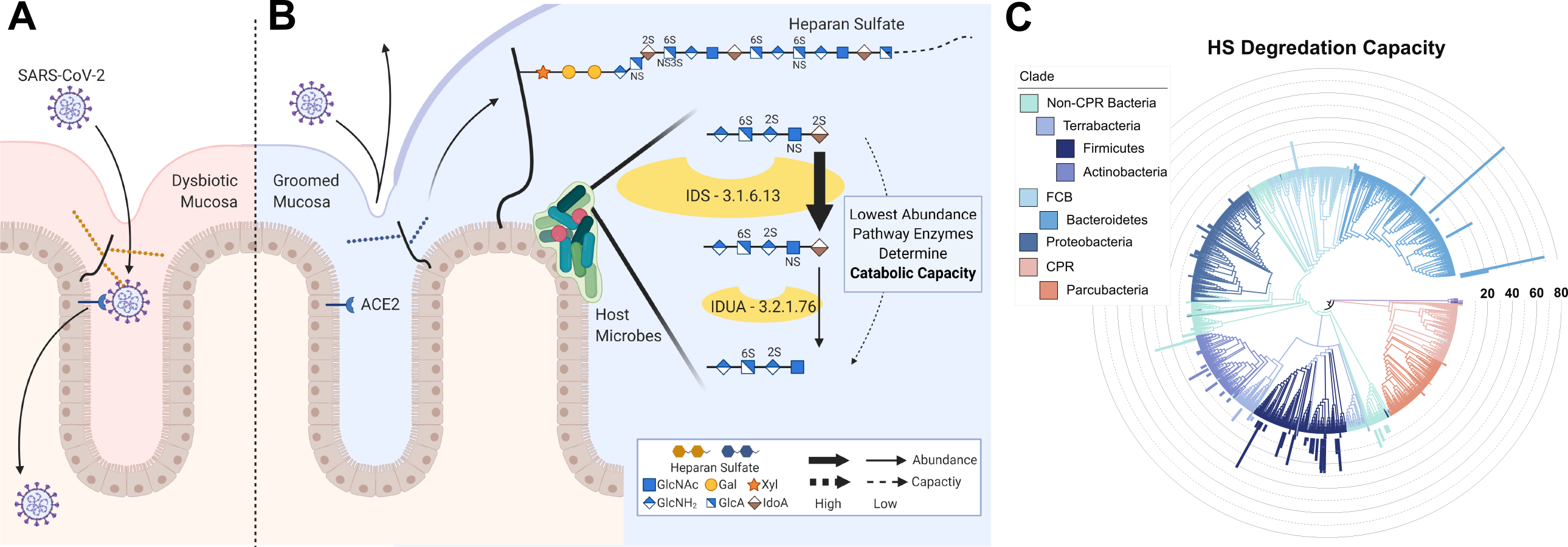
Prediction of bacterial capacity for modification of heparan sulfate (HS). Multiple studies have demonstrated microbial grooming of mucosal surfaces while separate studies have demonstrated the reliance of viruses on mucosal quality and quantity. Recently reliance on specific forms of HS, a major mucosal component, was demonstrated in SARS-CoV-2 (**A**). Here, we examine the abundance of HS-modifying genes in microbes measured in the FINRISK 2002 dataset and aggregate across the modification pathway to quantify the HS catabolic capacity of abundant microbes (**B**). Bacterial tree of life colored by superphylum groups and phyla containing predicted HS-modifying species. The bar chart represents the predicted capacity for HS modification of each species colored by phyla (**C**).

These counts allowed us to calculate HS catabolic “capacity” (a continuous measure of aggregate gene abundance within a pathway to estimate the amount of flux an organism can pass through the pathway) and “completeness” (binary indication of the presence or absence of all HS-catabolism genes in a microbe) (**Fig. 1C**). Specific inclusion criteria for HS modification genes and HS catabolic measures are described in **Materials and Methods**. Of the 7,490 species identified in the FINRISK 2002 dataset 463 species of bacteria, which we refer to as HS-modifying bacteria, contained the complete HS catabolic (completeness=1) task with high HS catabolic capacity (capacity>90^th^ percentile). Three *Bacteroides* species, *B. xylanisolvens, B. thetaiotaomicron* and *B. vulgatus*, were predicted to have extremely high capacity (99^th^ percentile) for HS catabolism consistent with prior reports (Cartmell et al., 2017; Ndeh et al., 2020; Ulmer et al., 2014).

### HS-modifying bacteria are reduced in COVID-19 patients

Based on the prevalence of bacteria that can potentially catabolize HS, we hypothesized that their presence within the microbiome might naturally limit HS-binding sites for SARS-CoV-2 and that the prevalence of bacteria encoding these genes might associate negatively with COVID-19 susceptibility (**Graphical Abstract**). To test this hypothesis, we explored the prevalence of predicted HS-modifying bacteria in COVID-19 patient samples. Human and SARS-CoV-2 reads were filtered out of previously published RNA-Seq datasets from 33 bronchoalveolar lavage fluid (BALF) samples from adult COVID-19 (n=13) and healthy control (n=20) subjects (Shen et al. 2019; Zhou et al. 2020) to obtain microbial reads (Poore et al., 2020). Gene expression from HS-modifying bacteria (N-strains numerator = 239, denominator = 1851) was decreased in COVID-19 compared to healthy control samples (**Fig. 2A**). Microbial RNA-Seq reads coding for heparin lyase (class I, II, & III) (**Fig. 2B)**, and N-acetylglucosamine-6-*O*-sulfatase (**Fig. 2C**), both HS-modifying enzymes, were also reduced in COVID-19 patients compared to healthy controls. These differences were not explained by variation between the two datasets included (**Fig. S1**). These findings suggest that COVID-19 patients have altered lung microbiomes that may favor SARS-CoV-2 infection.

**Figure 2.**
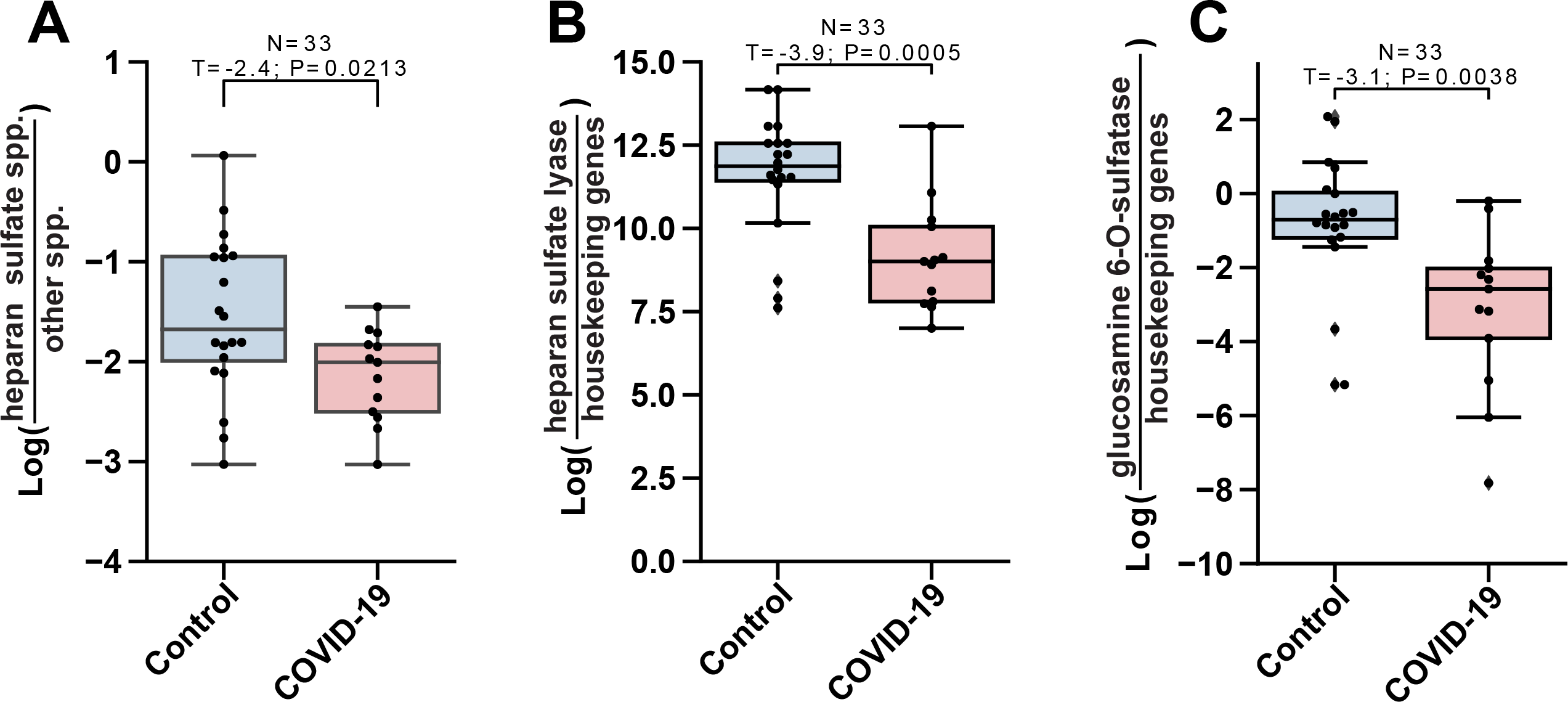
Bacteria species encoding heparan sulfate (HS) lyase (HSase) are depleted in COVID patients compared to controls. BALF RNA-seq data from healthy subjects (control) and COVID-19 patients (COVID) (x-axes) compared by log-ratios (y-axes) of predicted HS-modifying species relative to non-HS-modifying (**A**), HSase relative to housekeeping set (**B**), and N-acetylglucosamine-6-O-sulfatase relative to housekeeping set (**C**). Significance was evaluated by a t-test and error bars represent the standard error of the mean. Presented p-values are from unpaired two-tailed t-test.

### Bacteria with capacity for HS modification decrease with age

To determine if HS-modifying bacteria could reduce the risk of SARS-CoV-2 infection, we explored their prevalence across age and sex, two established COVID-19 risk-factors. Strong age and sex disparities in SARS-CoV-2 infection susceptibility has been reported across numerous populations, with adults twice as likely to be infected as those under the age of 20 (Davies et al., 2020) and men having higher risk than females (Williamson et al., 2020). Microbial communities age naturally and predictably with their human hosts in a sex-dependent manner (McDonald et al., 2018; de la Cuesta-Zuluaga et al., 2019; Claesson et al., 2011; Huang et al., 2020; Koenig et al., 2011), suggesting that aging and sex may be associated with a reduction in HS-modifying bacteria.

To test this hypothesis, we measured the prevalence of HS-modifying bacteria across age for both men and women in the FINRISK 2002 study. The log-ratio of bacteria predicted to catabolize HS versus all other bacteria (N-strains numerator = 687, denominator = 5987) decreased with age and in men compared to women as did the normalized abundance of bacterial HS-modifying genes (**Fig. 3A-B, Table S1)**. This finding was validated in an independent dataset by mapping the Web of Life (WoL) microbial genomes (Zhu et al., 2019) to 16S rRNA gene amplicon sequence variants (ASVs) from over 20,000 samples in the American Gut Project (AGP), a citizen-science dataset with participants ranging in age from 1-80 years (McDonald et al., 2018). A similar age and sex-dependent decrease in HS-modifying bacteria was observed (N-strains numerator = 244 & denominator = 1605) (**Fig. 3C**).

**Figure 3.**
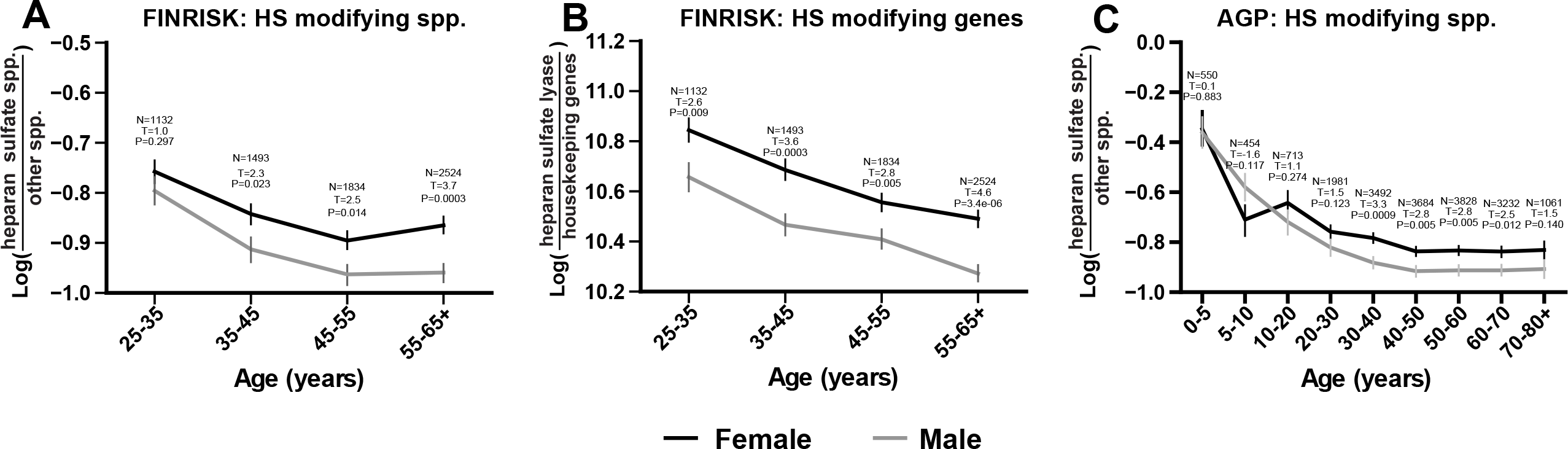
HS-modifying bacteria are depleted with host age and sex. The log-ratio of HS lyase genes relative to a set of housekeeping genes (y-axis) in the FINRISK 2002 fecal data compared by host age (x-axis) (**B**). The log-ratio of predicted HS-modifying species relative to those with no predicted capacity for (y-axes) in the FINRISK 2002 dataset (**A**), and AGP fecal dataset (**C**) compared over host age (x-axes). Log-ratios are colored by participant sex being female (black) and male (gray). All log-ratio plots annotated by the number of subjects at that time point. Error bars represent the standard error of the mean. Presented p-values and test statistics are from unpaired two-tailed t-test evaluated on each host age group between host sex.

Additional risk factors for COVID-19 identified in the OpenSAFELY study (Williamson et al., 2020) including diabetes, smoking, BMI, cancer, asthma, liver fibrosis or cirrhosis, cardiovascular, and autoimmune diseases were not observed to have significant changes by HS-modifying bacterial abundance in the FINRISK 2002 dataset (**Table S1**). The alpha diversity of both the AGP and FINRISK datasets exhibited a previously observed plateau after the age of 40 and an increase in women over men between the ages of 30 to 60 (de la Cuesta-Zuluaga et al., 2019; **Fig. S2A&B**). Alpha diversity and HS-modifying bacteria had a weak negative correlation for both men and women in the AGP and FINRISK datasets (**Table S2**).

### Common human gut commensal bacteria are capable of degrading HS

To provide more direct evidence for our hypothesis, *in vitro* experiments were conducted to test the catabolic capacity of human microbiome species to degrade HS. Axenic cultures of *B. ovatus* and *B. thetaiotaomicron*, highly prevalent human gut bacterial isolates known to catabolize HS (Cartmell et al., 2017) found in 80% of AGP and 99% of FIRNISK 2002 participants, were grown on minimal medium in the presence of heparin (100 µM) with and without glucose (22 mM). Both species were able to grow with heparin as the sole carbon source and electron donor (**Fig. 4A-B**). Furthermore, all cultures were verified to catabolize heparin in culture by comparing the concentration of heparin in the medium before and after stationary growth (**Fig. 4C-D**). Cell-surface HS on H1299 human lung cells was reduced by 60% upon exposure to cell-free supernatants from mid-log phase cultures of *B. ovatus* or *B. thetaiotaomicron* compared to 100% reduction by purified heparin lyase from *Flavobacterium heparinum* (*F. hep*. HSase; IBEX pharmaceuticals), as measured by binding of the anti-HS monoclonal antibody 10E4 (**Fig. 4E**).

**Figure 4.**
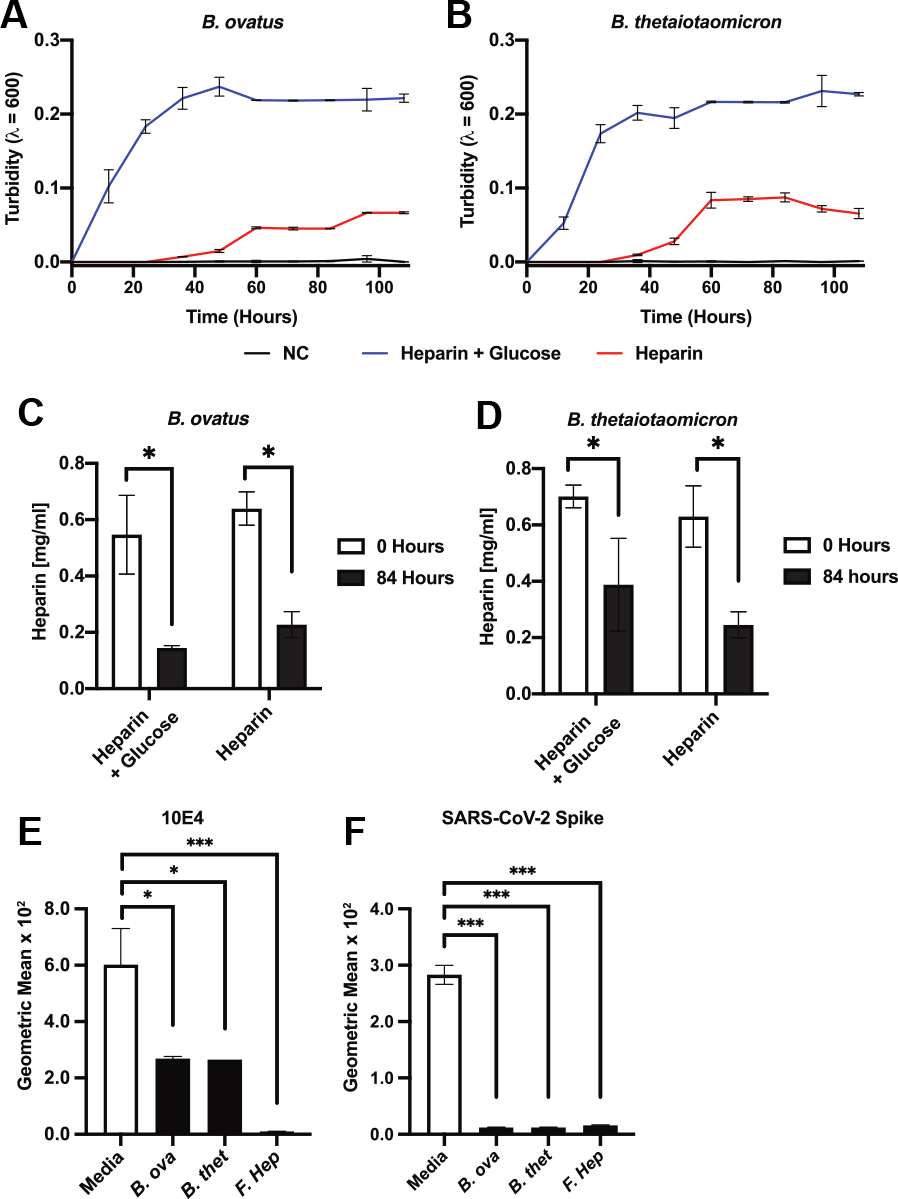
HS-modifying bacteria degrade HS and reduce SARS-CoV-2 spike protein binding when applied to human lung cells. Growth of *Bacteroides ovatus* (**A**) and *Bacteroides thetaiotaomicron* (**B**) measured by optical density (y-axis) across time from inoculation (x-axis) in minimal media (black; negative control NC), minimal media with 22 mM glucose and 100 µM heparin (blue), or minimal media with 100 µM heparin (red). Comparison of heparin concentration (y-axis; mg/ml) before inoculation (white; zero hours) and at stationary phase (gray; 84 hours) for *B. ovatus* (**C**) and *B. thetaiotaomicron* (**D**). Geometric mean of FACS count data (y-axis) of H1299 cells stained with the HS antibody 10E4 (**E**) or incubated with biotinylated SARS-CoV-2 spike protein (**F**) when untreated (NC), or incubated with culture media negative control (Media), cell-free supernatant of *B. ovatus* (B. ova) or *B. thetaiotaomicron* (*B. thet*.) or purified HSase from *Flavobacterium heparinum* (*F. hep*.). Presented p-values are from unpaired t-test statistics (p > 0.05 [n.s., not significant], p ≤ 0.05 [*], p ≤ 0.01 [**], p ≤ 0.001 [***]).

To determine the effect of bacterial HS modification on SARS-CoV-2 binding, H1299 human lung cells were treated with the supernatant of *Bacteroides* cultures or purified *F. hep*. HSase, and then incubated with biotinylated trimeric SARS-CoV-2 spike protein and assessed cell-surface binding by flow cytometry. Cells treated with *Bacteroides* culture supernatant caused a significant reduction (20-30 fold) in SARS-CoV-2 spike protein binding to cells incubated with *Bacteroides* culture supernatants compared to untreated H1299 cells (*B. ovatus* t-statistic = 31.25, p-value = 7.019 × 10^−5^; *B. thetaiotaomicron* t-statistic = 30.99, p-value = 7.023 ⨯ 10^−5^), similar to the reduction observed by pre-treatment with purified Hsase (t-statistic = 23.89, p-value = 7.59 ⨯ 10^−5^) (**Fig. 4F**).

## Discussion

We provide evidence for a role of the human microbiome in mediating SARS-CoV-2 infectivity via modification of the host glycocalyx. HS-modifying bacteria, identified in the human gut microbiome, were depleted in BALF of COVID-19 patients. Although no protective mechanism has been demonstrated yet, HS-modifying *Bacteroides* strains have previously been shown to be more frequent in healthy controls compared to COVID-19 patients (Trottein and Sokol, 2020; Zuo et al., 2020). *B. ovatus* and *B. thetaiotaomicron* can catabolize cell-surface HS and block SARS-CoV-2 S protein binding. In two independent gut microbiome datasets, we observed reduced HS-modifying bacteria, including *Bacteroides*, with increasing age and in men compared to women, which may help explain increased susceptibility to COVID-19 among men and older populations. Finally, we showed experimentally that *B. ovatus* and *B. thetaiotaomicron* can catabolize cell-surface HS and block SARS-CoV-2 S protein binding.

Human-associated microbes, including the *Bacteroides* species tested here, may alter viral infectivity in other ways. For example, ACE2, the plasma membrane receptor necessary for viral uptake, was downregulated in germ-free mice mono-colonized by *B. ovatus* or *B. thetaiotaomicron* (**Fig. S3**, Geva-Zatorsky et al., 2017). *Bacteroides* species, including *B. ovatus* and *B. thetaiotaomicron*, have previously been found to be negatively associated with host age (Galkin et al., 2020). Recent work has demonstrated that the receptor binding domain of SARS-CoV-2, unlike previous SARS-like coronaviruses, must be pre-activated by the proprotein convertase furin to effectively bind ACE2 (Walls et al., 2020, Hoffman 2020). Certain bacteria can also produce furin-like proteases and may provide another layer of regulation on top of host-produced furin proteases (Pavlova 2019). More generally, the effect of the microbiome on immune system development and decay has been widely documented (Thaiss et al., 2016), and may also influence an individual’s susceptibility to COVID-19.

We observed in the FINRISK 2002 dataset that non-modifiable COVID-19 predisposing factors such as host age and sex exhibit differences by bacterial HS-modification. Other risk factors such as diabetes identified in the OpenSAFELY project (Williamson et al., 2020) showed no clear trends by HS-modification. This highlights a possible critical role for microbial modulation in populations with non-modifiable risk factors, while other factors may be more accessibly modified by treating underlying metabolic imbalances. Furthermore, microbial communities obtained from other body sites such as saliva may also alter viral susceptibility in similar ways. For example, the human oral microbiome from the AGP dataset demonstrated an age dependent change in HS-modifying bacteria similar to the gut microbiome samples (**Fig. S4**). However, there are not comparably large oral microbiome datasets which could be used to explore other risk factors.

Positioned at the cell-environment interface, glycosaminoglycans (GAG), like HS, play critical roles in intercellular and host-pathogen interactions ranging from peptide exchange to migration (Weiss et al., 2017). Removal of critical molecules like HS may mitigate SARS-CoV-2 infection, but their depletion could also result in the loss of its endogenous function, for example in signaling reactions and tissue repair processes. Further studies are required to determine the target sulfation pattern for SARS-CoV-2 binding, but our results suggest that N-acetylglucosamine-6-*O*-sulfatase is decreased in the microbial populations present in COVID-19 patients. Activation of host 6-O-sulfotransferases decreased spike protein binding and dramatically reduced SARS-CoV-2 infection in tissue culture (Clausen and Sandoval et al., 2020), suggesting that 6-*O*-sulfation of HS may provide a more specific target to disrupt the SARS-CoV-2/HS interaction without broad degradation of host cell surface or extracellular matrix HS.

This report provides evidence for the role of glycocalyx-microbiota interactions as a novel competitive mechanism in the fight against SARS-CoV-2. Additional work will clarify and quantify the translational implications for microbial modulation of host GAGs and SARS-CoV-2 infectivity. It remains unclear which specific microbial enzymes modify GAGs to interrupt SARS-CoV-2 binding, and whether these enzymes can be exploited to limit the spread of the virus during an active infection. Additionally, the role of the respiratory tract microbiome, which has a strikingly different microbial community and functional repertoire from the gut microbiome (Human Microbiome Project Consortium, 2012; Beck et al., 2015; Lloyd-Price et al., 2017), remains to be explored. The analyses presented here describe a key mechanism by which host microbiomes might play a role in SARS-CoV-2 infection, provide a potential explanation for health disparities among age groups in COVID-19 disease and highlight the importance of characterizing the potential future molecular arms race between humans and viruses. Exploration of the vast interplay between host-microbes and viral infection could allow for improved risk stratification, prevention, and intervention.

## Methods

### Heparan Sulfate Modification Capability: Completeness and Capacity

The HS modification gene set is a metabolic task, i.e. a group of reactions necessary to transition between metabolites (Richelle et al., 2020) used to describe a complete pathway of HS modification. Pathways from the Kyoto Encyclopedia of Genes and Genomes (KEGG, ec00531, (Kanehisa et al., 2017)), in combination with literature annotation of these pathways for specific bacteria (Hobbs et al., 2018; Robb et al., 2017; Ndeh et al., 2017), were used to identify reactions (enzyme commission (EC) numbers) associated with the modification task. (**Table S3**) Finally, we used CAZy, dbCAN, and CUPP (Barrett and Lange, 2019; Lombard et al., 2014; Yin et al., 2012; Zhang et al., 2018) to map EC numbers to microbial genes associated with glycan modification (**Table S3)**.

We created two metrics to describe the HS-catabolic capabilities of microbes. For each microbe in the FINRISK2002 dataset, we estimated pathway “Completeness,” a binary indication of the presence or absence of all genes in the task, and pathway “Capacity,” a continuous indication of the magnitude of flux that could travel through the task. Expression of genes associated with glycan catabolism were quantified by the total number of reads (total count) mapped between the HS-modification gene set and FINRISK 2002 sample shotgun data through Bowtie2. A pseudocount was added to each count and log-transformed, *log(count+1)*, to stabilize variance.

HS-catabolic completeness (*E*_*i*_) for each organism, *i*, with gene, *g*, is an indication that all reactions in a task are represented in an organism. Specifically, for each reaction (EC) in the HS task, we sum the log-counts aligned to glycosylation-related genes within an organism associated with that EC. If the sum of log counts aligned to an EC exceeds a threshold, *t*, the EC was marked as active in that organism. The threshold, *t*, was set at the 5th percentile of log-counts aligned to glycosylation-associated genes in the FINRISK 2002 database, 4.522 log-counts. If all ECs in the HS task were active in an organism, the HS task is considered complete.

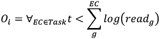

To quantify the HS-catabolic capacity (*A*_*i*_) of each organism, *i*, with genes, *g*, we analyzed the expression of all glycosylation-related genes within the catabolic task. Adapted from metabolic task analysis (Richelle et al., 2018, 2019, 2020), the catabolic capacity of a microbe is a context-sensitive measure that accommodates rate-limiting or low-abundance enzymes and can therefore accommodate microbial gene transience. Activation of the glycan degrading metabolic task (Richelle et al., 2020), was calculated as the minimum EC activation; the maximum activation of genes performing the EC reaction within an organism. Log of total counts was used to stabilize variance. The capacity is the minimum EC activation in a pathway where EC activation is the maximum gene activity score for each EC within a microbe. The log of total counts aligned to each gene in the FINRISK 2002 dataset was used for the gene activity score:

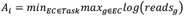

### FINRISK

Patient recruitment, sample collection, storage and molecular processing are described in detail in the flagship publication (Salosensaari et al., 2020). Demultiplexed shallow shotgun metagenomic sequences were quality filtered and adapter trimmed using Atropos (Didion et al., 2017), and human filtered using Bowtie2 (Langmead and Salzberg, 2012). For taxonomic assignment reads were aligned to the Web of Life database (Zhu et al., 2019) of 10,575 bacterial and archaeal genomes using Shogun (Hillmann et al., 2018) in the Bowtie2 alignment mode. For functional assignment of glycosylation-associated genes the FINRISK 2002 data were aligned to the HS degradation gene set of glycosylation-associated genes are using Bowtie2.

### Abundance and expression by log-ratios

Differential abundance or expression of HS-modifying bacteria was determined by the log-ratio of mapped reads for each sample of those bacteria with predicted capacity for HS modification relative to those without predicted capacity. Functional gene abundance or expression was determined with a log-ratio of the total mapped reads to that gene relative to the sum of counts of total mapped reads from a set of housekeeping genes in each sample. The housekeeping gene set is comprised of all bacterial nucleotide sequences for the genes atpD, dnaJ, gyrA, gyrB, infB, pheS, proC, rpoA, rpoB, and rpoD obtained from RefSeq (Pruitt et al., 2007). Significance between groups of log-ratios was determined with an unpaired t-test though SciPy (Virtanen et al., 2020).

### COVID-19 RNA-Seq dataset

Previously, COVID-19 metatranscriptomes have been analyzed for microbial taxonomy and function (Shen et al. 2020; Zhou et al. 2020; Haiminen et al. 2020). RNA-Seq data from two separate studies on SARS-CoV-2 infected patients (Shen et al., 2020; Zhou et al., 2020) were downloaded from the National Genomics Data Center (Project Accession PRJCA002202) and NCBI’s Sequence Read Archive (Project Accession PRJNA605983). Human and SARS-CoV-2 reads were filtered out by aligning reads to the human genome (Genome Reference Consortium Human Build 38) (Church et al., 2011; Schneider et al., 2017) and SARS-CoV-2 genome using Bowtie2 (Langmead and Salzberg, 2012) with the fast-local parameter set, following the protocol described by Poore et al. In the case of paired-end reads, forward and reverse reads files were treated as unpaired. Reads were aligned to the Web of Life database of 10,575 bacterial and archaeal genomes using Shogun (Hillmann et al., 2018) in the Bowtie2 alignment mode. Alignments were summarized into taxonomic profiles at the phylum, genus, and species level using the classify function in the Web of Life (Zhu et al., 2019).

### Mapping of WoL to AGP 16S ASVs

To reconcile the evolutionary relationships among 16S rRNA gene sequencing amplicon sequence variants (ASV) and shotgun metagenomic data, we mapped the ASVs to the “Web of Life” (WoL) (Zhu et al., 2019) reference phylogeny of bacterial and archaeal genomes. First, 16S rRNA genes were annotated from each of the 10,575 genomes included in the phylogeny using RNAmmer 1.2 (Lagesen et al., 2007), using domain-specific models (bacteria and archaea, respectively). Second, filtered 150 bp length 16S V4 American Gut Project (McDonald et al., 2018) ASVs (*n*=15,486) were aligned to the WoL 16S rRNA genes using BLASTn 2.7.1+ (Altschul et al., 1990), with an e-value threshold of 1e-5 and up to 100 target sequences per query. Top hits with identical bit scores of each query were retained and subjected to taxonomic classification. At each designated rank, taxonomic assignments of all top hits were recorded. For feature table generation, the hits were counted and normalized by the total number of hits. As an example, assuming one ASV aligned equally well to five reference full-length 16S sequences, and they belong to genus A (two sequences) and genus B (three sequences), then the two genera were counted as 2/5 (A) and 3/5 (B), respectively. Per-query counts were summed across each AGP sample and rounded to integers.

### Cultivation of Bacteroides strains

*Bacteroides thetaiotaomicron* and *Bacteroides ovatus* were cultured in 50 mL anaerobic brain heart infusion (BHI) medium with overnight incubation at 37°C in an anaerobic serum bottle. The next day, 1 mL of cells from the BHI culture were washed in anaerobic PBS and were passed into 15mL of minimal medium containing no carbon sources or electron donors other than glucose (22 mM final) and/or heparin (1 mg/mL final). Growth on the minimal medium was measured by optical density at 600 nm every 12 hr. Aliquots of 3 mL of culture were taken before inoculation, immediately after inoculation, at mid-log phase (24 hr), and in stationary phase (45 hr). Heparin degradation in culture was measured through a Blyscan glycosaminoglycan assay (Biocolor Ltd., Carrickfergus, Northern Ireland) using 100 µL of time 0- and 45-hr culture media. All cultivation and HS-modification experiments were conducted in triplicate.

To verify strain taxonomy a 1 mL aliquot of each culture was extracted using PowerFecal DNA Isolation Kit (MoBio cat. 12830). Extracted DNA was quantified via Qubit™ dsDNA HS Assay (Thermo Fisher Scientific), and 5 ng of input DNA was used in a 1:10 miniaturized Kapa HyperPlus protocol (Sanders et al., 2019). The pooled library was sequenced as a paired-end 150-cycle run on an Illumina NovaSeq at the UCSD IGM Genomics Center. Resulting sequences were adapter trimmed using Trimmomatic v0.39 (Bolger et al., 2014) and human read filtered with Bowtie2 (Langmead and Salzberg, 2012). Paired-end reads were merged using Flash v1.2.11 (Magoč and Salzberg, 2011). Each axenic sample was assembled through SPAdes (Bankevich et al., 2012) and verified through average nucleotide identity (Lee et al., 2016) of greater than 99% between the assembled genome and the putative type-strain genome obtained from NCBI (Pruitt et al., 2007).

### SARS-CoV-2 spike protein production

Recombinant SARS-CoV-2 spike protein encoding residues 1-1138 (Wuhan-Hu-1; GenBank: MN908947.3) with proline substitutions at amino acids positions 986 and 987 and a “GSAS” substitution at the furin cleavage site (amino acids 682-682), was produced in ExpiCHO cells by transfection of 6 x106 cells/ml at 37 °C with 0.8 □g/ml of plasmid DNA using the ExpiCHO expression system transfection kit in ExpiCHO Expression Medium (ThermoFisher). One day later the cells were refed, then incubated at 32 °C for 11 days. The conditioned medium was mixed with cOmplete EDTA-free Protease Inhibitor (Roche). Recombinant protein was purified by chromatography on a Ni2+ Sepharose 6 Fast Flow column (1 ml, GE LifeSciences). Samples were loaded in ExpiCHO Expression Medium supplemented with 30 mM imidazole, washed in a 20 mM Tris-Cl buffer (pH 7.4) containing 30 mM imidazole and 0.5 M NaCl. Recombinant protein was eluted with buffer containing 0.5 M NaCl and 0.3 M imidazole. The protein was further purified by size exclusion chromatography (HiLoad 16/60 Superdex 200, prep grade. GE LifeSciences) in 20 mM HEPES buffer (pH 7.4) containing 0.2 M NaCl.

### Biotinylation

For binding studies, recombinant SARS-CoV-2 spike protein was conjugated with EZ-Link™ Sulfo-NHS-Biotin (1:3 molar ratio; Thermo Fisher) in Dulbecco’s PBS at room temperature for 30 min. Glycine (0.1 M) was added to quench the reaction and the buffer was exchanged for PBS using a Zeba spin column (Thermo Fisher).

### SARS-CoV-2 spike binding experiments

NCI-H1299 from the American Type Culture Collection were grown in RPMI medium containing 10% FBS and 100 U/mL penicillin and 100 µg/mL streptomycin sulfate under an atmosphere of 5% CO_2_ and 95% air. Cells at 50-80% confluence were lifted in 10 mM EDTA in PBS (Gibco) and washed in PBS containing 0.5% BSA. Cells were seeded into a 96 well plate at 10^5^ cells per well. The cells were then treated with 2.5 mU/mL *Flavobacterium heparinum* HSase I, 2.5 mU/mL *Flavobacterium heparinum* HSase II, and 5mU/mL *Flavobacterium heparinum* HSase III (IBEX Pharmaceuticals) in PBS containing 0.5% BSA (100 µL) or 100 µL *Bacteroides ovatus* and *thetaiotaomicron* minimal media culture cell-free supernatant supplemented with 10% BSA) for 30 min at 37 °C. A portion of the cells was treated with HSase mix for 30 min at 37°C in PBS containing 0.5% BSA. The cells were washed twice in PBS containing 0.5% BSA. The cells were then stained with 20 µg/mL biotinylated S protein (S1/S2) or 1:1000 Anti-HS (Clone F58-10E4) (Fisher Scientific, NC1183789) in PBS containing 0.5% BSA, for 30 min at 4°C. The cells were washed twice and then stained with Streptavidin-Cy5 (Thermo Fisher) at 1:1000 (S protein) or Anti-IgM-Alexa488 at 1:1000 (Anti-HS) in PBS containing 0.5% BSA, for 30 min at 4°C. The cells were washed twice and then analyzed using a FACSCanto instrument (BD bioscience). Data analysis was performed in FlowJo and statistical analyses were done in Prism 8.

## Data availability

The Shen et al. data is available and Zhou et al. data is available through the National Genomics Data Center accession PRJCA002202 and NCBI’s Sequence Read Archive accession PRJNA605983 respectively. All American Gut Project sequence data and deidentified participant responses can be found in EBI under project PRJEB11419 and Qiita (https://qiita.ucsd.edu/) study ID 10317. The FINRISK data that support the findings of this study are available from the THL Biobank based on a written application and following relevant Finnish legislation. Details of the application process are described in the web-site of the Biobank: https://thl.fi/en/web/thl-biobank/for-researchers.

## Code availability

The source code for the analyses can be found at https://doi.org/10.5281/zenodo.3973506.

## Author Contributions

C.M., B.P.K, T.M.C., D.R.S., J.D.E., N.E.L., and R.K. conceived, initiated, and coordinated the project. B.P.K, E.A., J.S., C.L., Y.Z., and N.E.L. conceived and performed pathway models. C.M., T.M.C., D.R.S., L.Z., and R.A.S.B. designed and performed the experimental work. C.M., B.P.K, A.S., S.J.S., S.W., E.A., Y.V.B., D.M., L.Z., S.C., Q.Z., B.T., M.I., V.S., M.J., A.S.H., P.J., T.N., and R.K. coordinated, compiled and analyzed sequencing data. B.P.K, T.M.C., K.Z., P.B.F, L.S., N.R.K., J.D.E., and R.K. supplied reagents. C.M., B.P.K., T.M.C., D.R.S., C.A.M., J.D.E., N.E.L., and R.K. wrote the manuscript with input from all authors. All authors discussed the experimental results and read and approved the manuscript.

## Conflicts of Interest

VS has received honoraria for consultations from Novo Nordisk and Sanofi. He also has ongoing research collaboration with Bayer Ltd (all unrelated to the present study).

## Acknowledgements

This work was supported in part by IBM Research AI through the AI Horizons Network, NIH Pioneer award 1DP1AT010885, RAPID grant 2038509 from the National Science Foundation, University of California on COVID-19 grant R00RG2503, and Emerald Foundation 3022 (to R.K.), the NIDDK grant 1P30DK120515, NIGMS R35GM119850 (to N.E.L.), RAPID grant 2031989 from the National Science Foundation and Project 3 of NIH P01 HL131474 (to J.D.E.). The Alfred Benzon foundation (to T.M.C.). The Academy of Finland grant 321351 and the Emil Aaltonen Foundation (to T.N.). The National Institutes of Health grant R01ES027595 (to M.J.). The ANID Becas Chile Doctorado 2018-72190270 (E.A.). The Academy of Finland grants 321356 and 335525 (A.S.H)

## Supplemental Figures

**Supplemental Figure 1.**
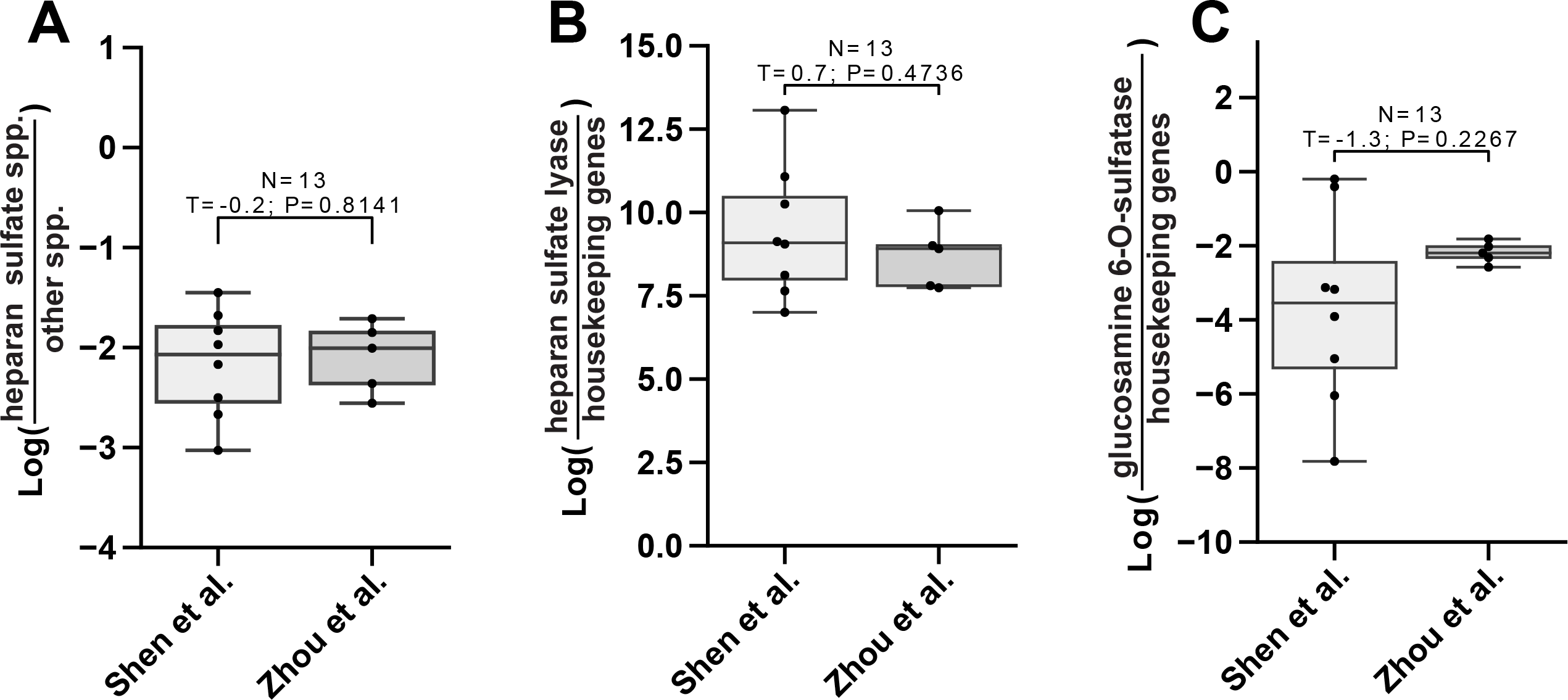
BALF RNA-seq data compared between studies. RNA-seq studies (x-axes) compared by log-ratios (y-axes) of predicted HS-modifying species relative to non-HS-modifying (**A**), HSase relative to housekeeping set (**B**), and N-acetylglucosamine-6-O-sulfatase relative to housekeeping set (**C**).

**Supplemental Figure 2.**
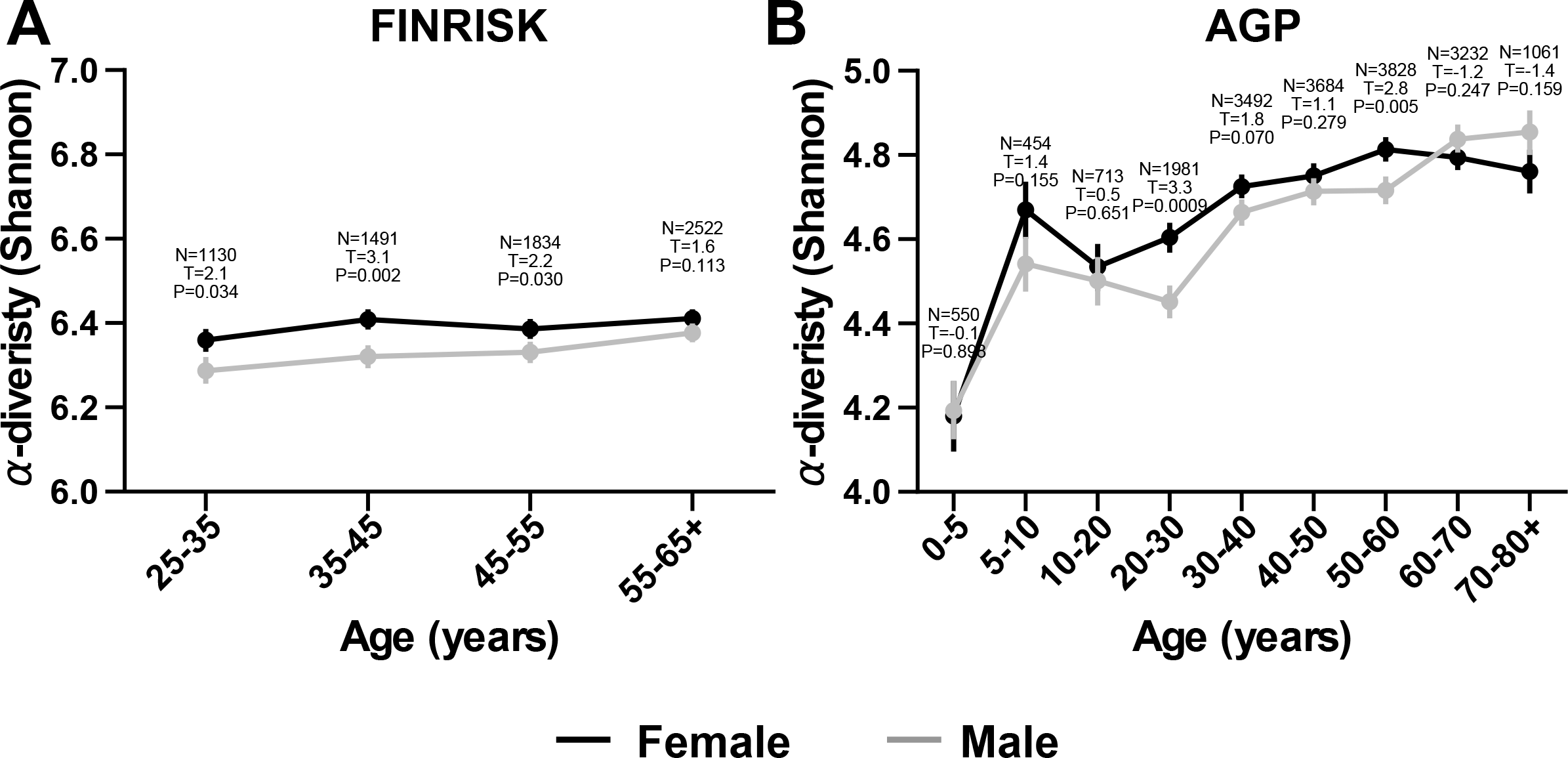
Shannon alpha diversity (y-axes) changes by age (x-axes) and colored by host sex being women (black) and men (gray) in FINRISK (A) and AGP (B) datasets.

**Supplemental Figure 3.**
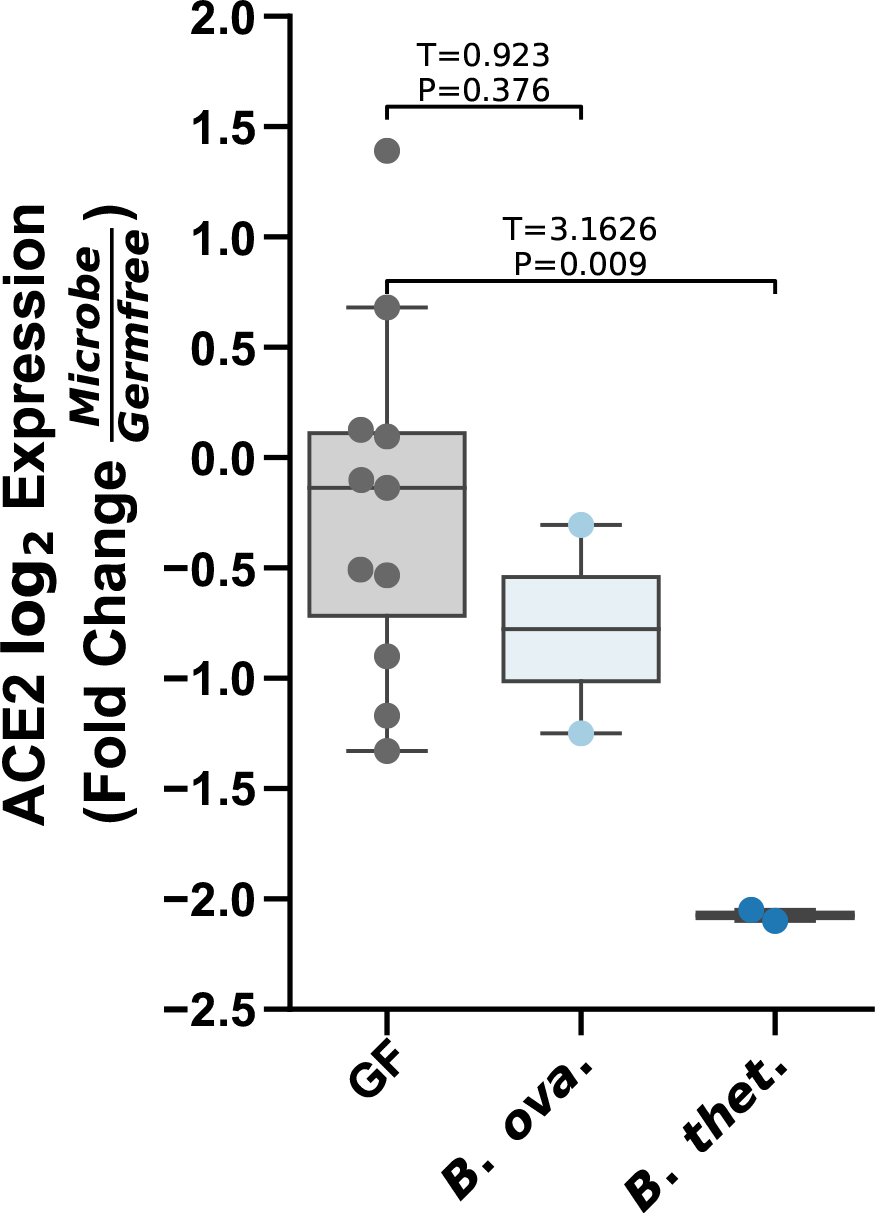
Bacteroides strain colonization associated with down regulated ACE2 in the colon relative to germ free (GF) mice. Presented p-values are from unpaired two-tailed t-test statistics.

**Supplemental Figure 4.**
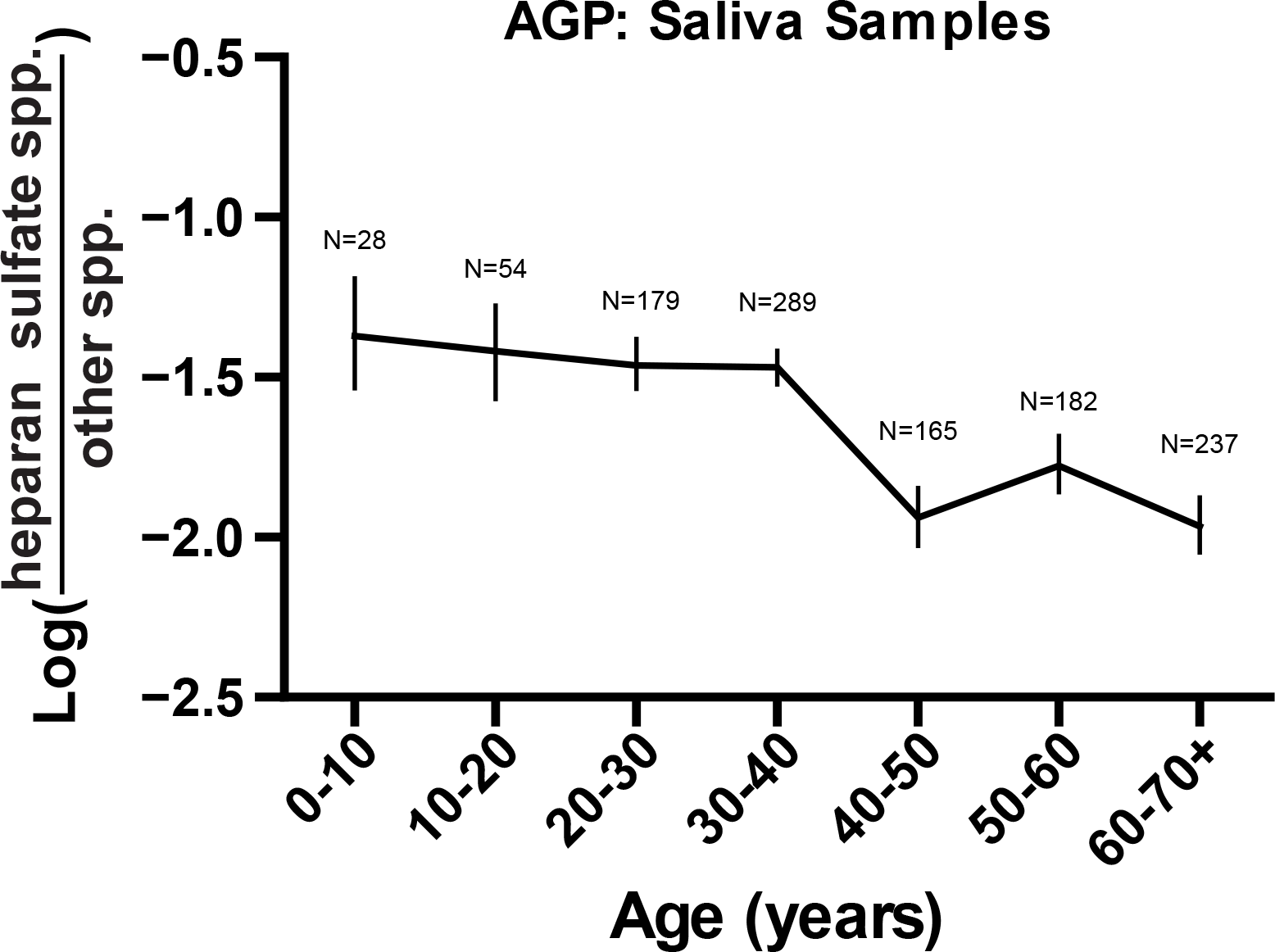
The log-ratio of predicted HS-modifying species relative to those with no predicted capacity for (y-axes) AGP saliva samples across age (x-axis).

## Supplemental Tables

**Supplemental Table 1**. COVID-19 risk factors in relation to the log-ratio of predicted HS-modifying species relative to those with no predicted capacity assed by OLS-regression in the FINRISK 2002 dataset.

**Supplemental Table 2**. Pearson correlation shows weak correlation between alpha diversity (Shannon) and heparan sulfate modifying log-ratio split by host sex.

**Supplemental Table 3**. Table of CAZy ID associated with heparan sulfate modification tasks for each bacterial species along with the capacity and completeness for a given species.

